# *C. elegans* ELT-3 regulates cuticle collagen expression in response to environmental stimuli

**DOI:** 10.1101/2020.03.06.980656

**Authors:** Hiva Mesbahi, Kim B. Pho, Andrea J. Tench, Victoria L. Leon Guerrero, Lesley T. MacNeil

## Abstract

The nematode *Caenorhabditis elegans* is protected from the environment by the cuticle, an extracellular collagen-based matrix that encloses the animal. Over 170 cuticular collagens are predicted in the *C. elegans* genome, but the role of each individual collagen is unclear. Stage-specific specialization of the cuticle explains the need for some collagens, however, the large number of collagens suggests that specialization of the cuticle may also occur in response to other environmental triggers. Missense mutations in many collagen genes can disrupt cuticle morphology, producing a helically twisted body causing the animal to move in a stereotypical pattern described as rolling. We find that environmental factors, including diet, early developmental arrest, and population density can differentially influence the penetrance of rolling in these mutants. These effects are in part due to changes in collagen gene expression that are mediated by the GATA family transcription factor ELT-3. We propose a model by which ELT-3 regulates collagen gene expression in response to environmental stimuli to promote the assembly of a cuticle specialized to a given environment.

## Introduction

The ability to respond to environmental cues is required for animals to survive in changing environments and allows an animal flexibility in the environments it inhabits. In the wild, *C. elegans* is exposed to frequent changes in environment and has developed strategies to cope with pathogens, temperature, osmolarity, and intermittent availability of oxygen or food. For *C. elegans*, the first barrier against environmental insults is the cuticle, which acts as a physical barrier to protect the animal from pathogens, desiccation, and other stresses (Cassada and Russell 1975; Gravato-Nobre *et al.* 2016; Xiong *et al.* 2017). The cuticle is a multi-layered, flexible, collagen-rich exoskeleton synthesized by underlying hypodermal cells (Cox *et al.* 1981). During development, it is shed at each molt to allow for growth and is resynthesized five times during the life of the animal. Although not all larval (L) stages have been examined in detail, L1, L4, and adult cuticles differ in morphology and collagen composition (Cox *et al.* 1981; Cox and Hirsh 1985). In the dauer state, an alternate larval stage specialized for long term survival and resistance to desiccation, the cuticle is thicker and structurally distinct from cuticles at all other stages (Popham and Webster 1978; Cox *et al.* 1981; Peixoto and De Souza 1994). Stage-specific cuticles are generated through transcriptional regulation of specific collagen genes. This regulation ensures that the appropriate collagens are expressed at the correct time and allows specialization of the cuticle for each larval stage (Cox *et al.* 1981; Cox and Hirsh 1985; Jackson *et al.* 2014; Teuscher *et al.* 2019).

Modulation of collagen expression may be used to promote resistance to specific stresses. In fact, the expression of several collagens is modulated in response to exposure to different bacterial species (Coolon *et al.* 2009), and environmental stresses, such as high salt (Dodd *et al.* 2018). The expression of specific collagens may promote increased survival in specific environments. Consistent with this idea, interventions that extend lifespan, and reduce insulin-like signaling, influence the expression of many collagen genes, suggesting that the ability to remodel the cuticle may contribute to lifespan extension (Ewald *et al.* 2015).

The *C. elegans* genome contains 181 predicted collagen genes, 173 of which are considered cuticular collagens (Teuscher *et al.* 2019). *C. elegans* collagens have been broadly classified into five groups based on the position of conserved cysteines (Johnstone 1994; Page and Johnstone 2007). Nematode collagens are most similar to fibril-associated collagens with interrupted triple helices but differ from vertebrate collagens in that they are generally smaller and less complex. These collagens assemble into trimeric collagen fibrils that are assembled before secretion. Collagen proteins are characterized by the presence of Gly-X-Y domains, where X and Y are enriched for proline and hydroxyproline, and several conserved cysteine residues. The Gly-X-Y domains are required for the formation of triple helical structures while the cysteines form intermolecular disulfide bridges. Both homo and heterotrimeric complexes have been described for vertebrate collagens and *C. elegans* collagens are likely to be organized in a similar manner.

Mutations in 21 cuticular collagen genes have been isolated from unbiased genetic screens (Cox *et al.* 1980; Page and Johnstone 2007). These mutations result in changes in body size and morphology, producing dumpy (Dpy) or long (Lon) animals, blistered animals, and rolling animals. The rolling movement observed in these mutants is the result of a twisted cuticle that causes the animal to rotate around its long axis and move in an abnormal stereotypical pattern. Mutations in several collagens, including *sqt-1*, *dpy-17*, *lon-3*, *rol-6*, *dpy-5*, and *bli-1* produce observable phenotypes, and different mutations in the same gene can produce distinct phenotypes. For example, different mutations in *sqt-1* can produce dumpy, long, or rolling animals (Kusch and Edgar 1986). Mutations that result in rolling are often dominant mutations, whereas null alleles of these genes do not produce rolling, suggesting that the rolling phenotype results from the incorporation of aberrant collagen molecules into the cuticle. For example, while dominant mutations in *rol-6* produce rollers, null alleles are superficially wild type (Kramer *et al.* 1990).

Mutations in *rol-6*, *sqt-3*, *dpy-10*, and *sqt-1* that produce a rolling phenotype have been isolated. These mutants are classified as right or left rollers depending on the direction of the roll (Cox *et al.* 1980). Mutations in *sqt-3* and *dpy-10* produce left rollers, whereas *rol-6* and *sqt-1* mutations produce right rollers (Cox *et al.* 1980). Intriguingly, rolling in these mutants can be suppressed by loss or knockdown of other collagens (Kramer *et al.* 1990; Nyström *et al.* 2002). For instance, rolling in dominant *rol-6* mutants can be suppressed by loss of *sqt-3* or *sqt-1* (Kramer *et al.* 1990). Similarly, rolling in *sqt-3* and *sqt-1* mutants can be suppressed by knocking down *rol-6* (Kramer and Johnson 1993; Cai *et al.* 2011). One proposed explanation for these findings is that these collagens form heteromeric complexes or a common substructure and reducing levels of specific collagens may reduce the incorporation of aberrant collagen proteins in the cuticle (Johnstone 1994). This model is consistent with the finding that the expression of collagen genes capable of mutating to produce roller phenotypes is highly coordinated, with their expression occurring in a short window before the molt (Hendriks *et al.* 2014).

The dominant *rol-6 (su1006)* allele is commonly used as a co-injection marker when generating transgenic animals because it produces a readily observable phenotype with little toxicity. The penetrance of rolling in these strains is often incomplete, is temperature sensitive, and differs between strains. Further, we observed that the penetrance within a strain is not robust, but is sensitive to growth conditions. Based on these observations, we hypothesized that the penetrance of rolling is sensitive to environmental stimuli. We find that the penetrance of the rolling phenotype in mutants of the collagens *rol-6* and *sqt-3* is differentially affected by bacterial diet, metabolic disruption, starvation, and population density. By examining the expression of collagen genes, we identify collagens whose expression is modulated in response to these environmental factors and identify *elt-3* as a regulator of these collagens. We find that in *elt-3* mutants, the effect of environmental factors on rolling is reversed and hypothesize that *elt-3* functions in environmental-response pathways to induce changes in collagen gene expression that ultimately function to tailor the cuticle to specific environments.

## Results

### Environment influences the penetrance of the rolling phenotype

To examine environmental influence on rolling, we first tested the effect of diet on the penetrance of rolling in strains carrying a dominant *rol-6(su1006)* transgene [hereafter referred to as *rol-6(su1006)*T]. When r*ol-6(su1006)*T strains with incomplete penetrance were fed a diet of *Comamonas aquatica* DA1877, the penetrance of rolling was dramatically decreased compared to animals fed *E. coli* OP50 (Figure 1A and 1B). This suppression was observed in several integrated transgenic strains with incomplete penetrance of the rolling phenotype, but not in strains with complete, or nearly complete, penetrance, including the MS438 (*irIs25*[*elt-2*∷NLS∷GFP∷lacZ + *rol-6(su1006)*]) transgenic strain and the endogenous mutant allele, *rol-6(e178)* (Figure 1B). To further examine the effects of *C. aquatica* on the cuticle, we measured cuticle permeability by staining animals with Hoechst dye 33258. Compared to animals fed *E. coli* OP50, exposure to *C. aquatica* increased permeability of the dye, resulting in a larger number of positive staining animals (Figure 1C). Nuclear staining in *rol-6(su1006)*T animals was predominantly in the head and tail and generally produced weakly staining nuclei. Collectively, these data suggest that *C. aquatica* exposure alters the cuticle in *rol-6(su1006)*T animals.

**Figure 1.**
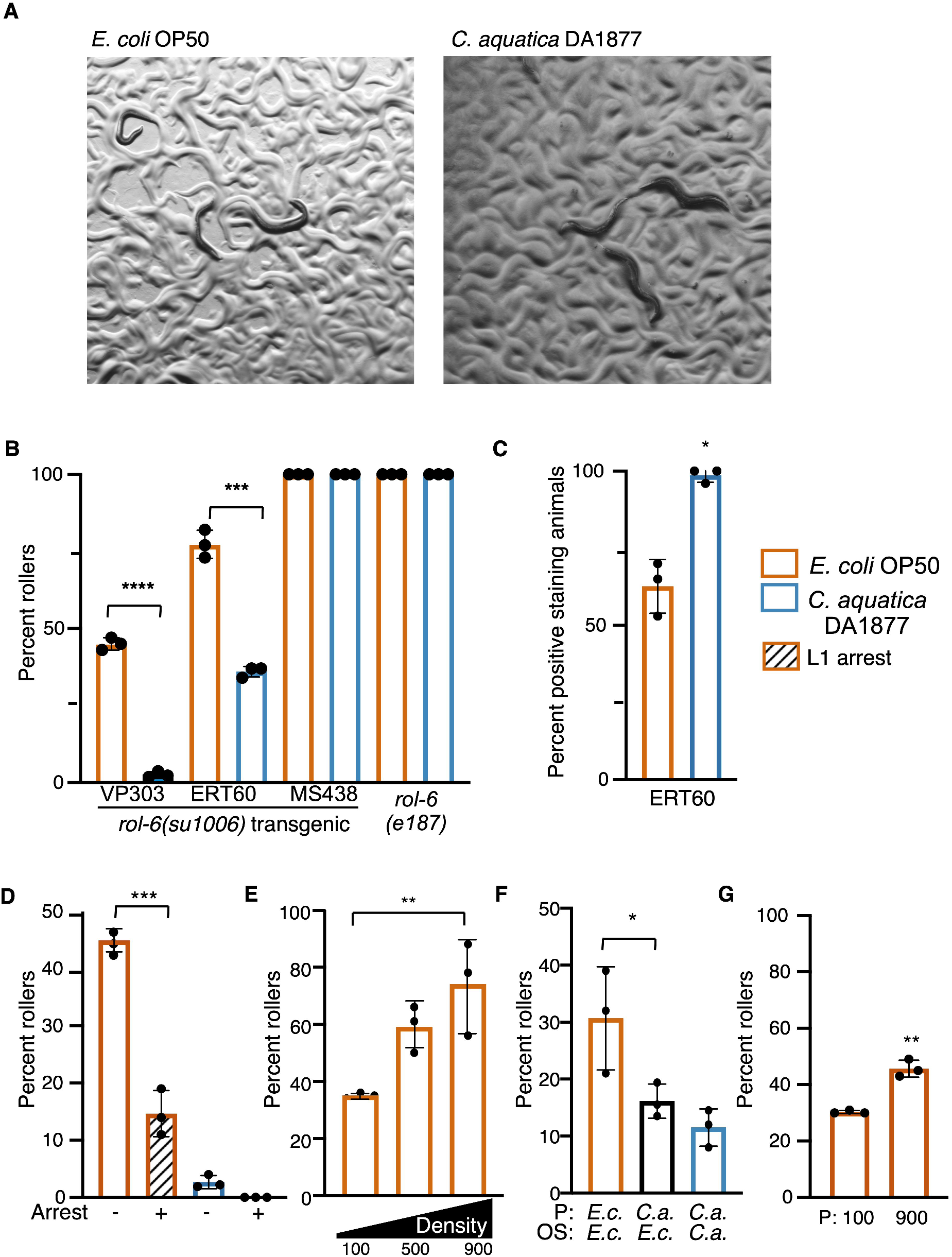
Environment influences rolling in animals carrying *rol-6* mutations. A. *rol-6(su1006)*T (VP303) animals exposed to *E. coli* OP50 and *C. aquatica* DA1877. B. Percent of population rolling when grown on indicated bacterial diets. Assays were done in triplicate; each dot represents a population of greater than or equal to 50 animals. C. Effect of *C. aquatica* exposure on rolling in three transgenic *rol-6(su1006)*T lines and in the mutant *rol-6(e187)* D. Hoechst staining of *rol-6(su1006)*T animals (ERT60) exposure to *E. coli* or *C. aquatica*. E. Percent rolling VP303 adults when offspring were treated by L1 arrest (+) or allowed to develop from eggs (−). F. Maternal exposure to *C. aquatica* reduces rolling in the next generation. Offspring (OS) from *E. coli* (E.c.) or *C. aquatica* (C.a.) fed parents (P) were scored (VP303). G. Percent of VP303 animals rolling at indicated densities (60mm dish). G. Intergenerational effects of population density in *rol-6(su1006)*T animals (VP303). Parental generation (P) was grown at high (900/60mm) or low density (100/60mm). B-E Statistical analysis by one way Anova and Dunnett’s multiple comparison test. F. Statistical analysis by one-way Anova with Tukey’s multiple comparison test. D and G statistical analysis using t-test. Asterisks indicate P-values (*<0.05, **<0.005, ***<0.0005).

To extend these findings, we examined the influence of two common environmental variables, starvation and population density, on the penetrance of rolling. To test the effect of starvation on rolling, we allowed L1 *rol-6(su1006)*T animals to undergo starvation-induced arrest and examined rolling in the adult. Relative to animals that had not undergone arrest, the penetrance of rolling was reduced when animals had previously undergone L1 arrest (Figure 1D). Next, we tested the effects of increased population density on rolling. Although starvation and high population density both represent unfavourable conditions for the worm (both promote dauer formation), and both could be viewed as conditions signaling limited resources, high population density produced the opposite effect on rolling as the L1 arrest. When 900 animals were present on a 60 mm plate, the incidence of rolling was approximately two-fold higher than when populations were 100 animals/plate (Figure 1E). Together, these observations suggest that many different environmental cues stimulate changes to the cuticle and that the integration of these cues may allow the animal to adapt to different environmental challenges.

### Environmental factors induce intergenerational effects

We asked whether environmental effects could be carried over to subsequent generations by testing whether parental exposure to *C. aquatica* could suppress rolling in the next generation. Eggs were collected by hypochlorite bleaching from animals grown on *C. aquatica*, or *E. coli* OP50, and offspring were developed on *E. coli* OP50. As adults, the offspring of *C. aquatica* fed parents display reduced rolling relative to those of *E. coli* OP50-fed parents (Figure 1F) suggesting that parental exposure can influence cuticle dynamics in the offspring. Similarly, we tested the effects of population density on rolling in the next generation. Eggs were collected from plates originally seeded with 100 or 900 animals and rolling was examined when the offspring became adults. Similar to what we observed with diet, parental population density influenced the penetrance of rolling, with more densely growing populations producing higher penetrance of rolling in the next generation (Figure 1G).

### Metabolic disruption influences rolling penetrance

*C. aquatica* and *E. coli* OP50 differ in the ability to produce vitamin B12 (Watson *et al.* 2014). *C. aquatica* produces vitamin B12, while *E. coli* does not. To determine whether vitamin B12 was responsible for the suppression of rolling in animals fed *C. aquatica*, we added B12 to the *E. coli* OP50 diet and measured rolling. Addition of vitamin B12 produced a dose-dependent decrease in rolling in *rol-6(su1006)*T strains with moderate to low penetrance rolling (VP303, NR222, FK181)(Figure 2A). However, similar to what was observed with diet, rolling in *rol-6(e187)* mutants was not suppressed by vitamin B12.

**Figure 2.**
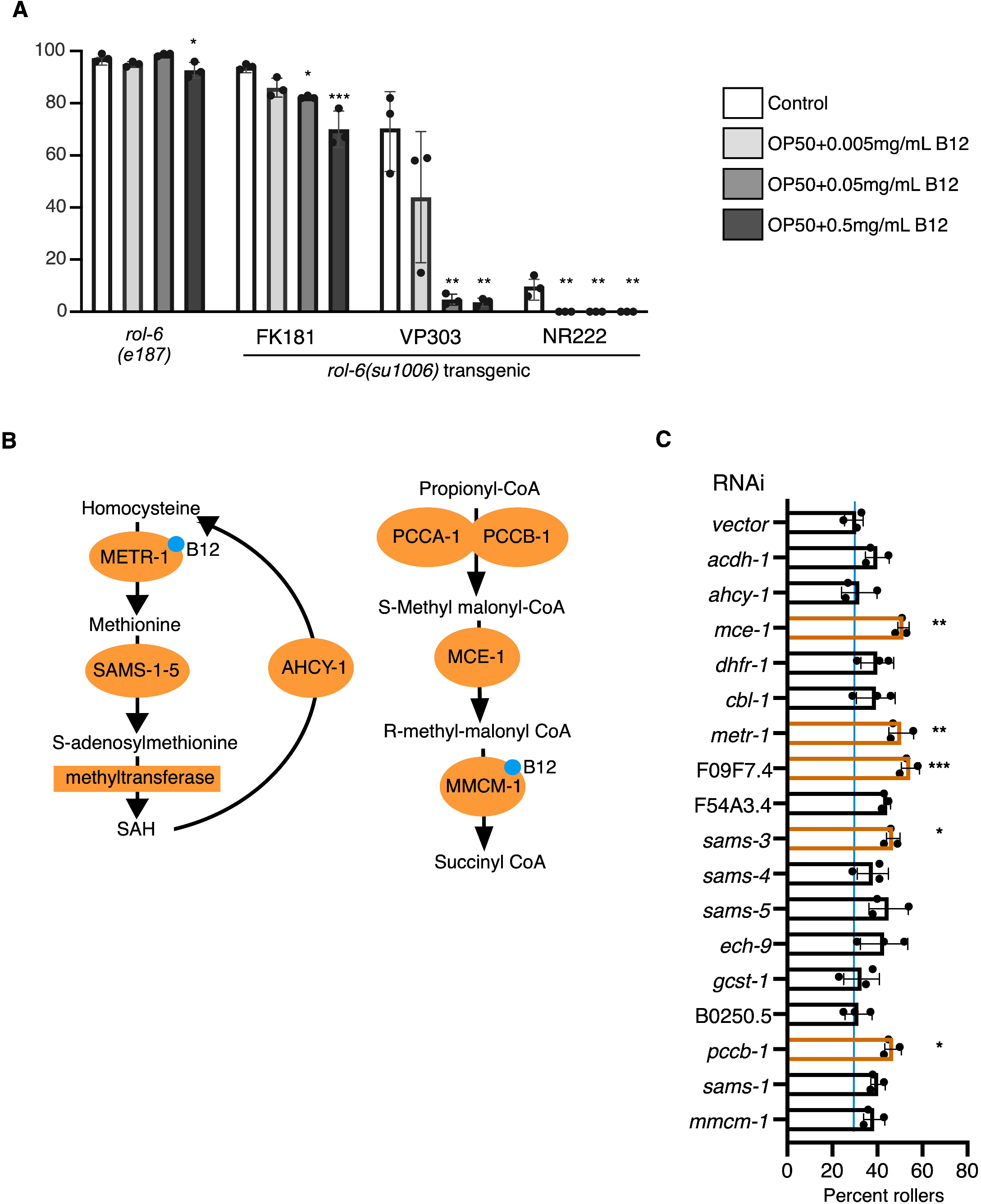
Metabolic disruption in the hypodermis influences rolling. A. Response of indicated animals to increasing concentrations of Vitamin B12. Vitamin B12 reduced rolling in *rol-6(su1006)*T animals. Significance refers to comparisons to controls of the same strain. B. B12 dependent metabolic pathways, enzymes that use vitamin B12 as a cofactor are indicated. C. RNAi-mediated knockdown of metabolic enzymes in B12-dependent pathways. RNAi was induced by feeding in a single generation (NR222, 25C). Percent rollers in a population is shown, each point represents a population of >50 animals. Bars represent mean, plus and minus the standard deviation. Statistical analysis by one-way Anova with Tukey’s multiple comparison test. Asterisks indicate P values (*<0.05, **<0.005, ***<0.0005).

Vitamin B12 acts as a cofactor for two enzymes, METR-1, functioning in 1-Carbon metabolism and methyl malonyl CoA mutase, MMCM-1, which converts methylmalonyl CoA to succinyl CoA (Figure 2B). To determine which activity was required for the observed suppression of rolling, we knocked down enzymes in both pathways and measured the effect on rolling behavior. Knockdown of *metr-1* produced the most dramatic effects, however, knockdown of enzymes in both pathways increased the penetrance of rolling in *rol-6(su1006)*T animals (Figure 2C), consistent with the finding that addition of vitamin B12 reduces rolling. These findings demonstrate that metabolic imbalances also have the potential to promote cuticle remodeling.

### Environmental effects on rolling penetrance are collagen-specific

Conceptually, suppression of rolling in *rol-6(su1006)*T animals could be mediated in two ways, through changes in gene expression that reduce the contribution of ROL-6 to the cuticle or by generating a robust cuticle more tolerant of the aberrant ROL-6 protein. If the latter were true, conditions that suppress rolling in *rol-6(su1006)*T animals might also suppress rolling resulting from mutation in other collagen genes. To test this hypothesis, we exposed animals carrying rolling-inducing alleles of *sqt-3, dpy-10, rol-1 and rol-9* to conditions that suppress rolling in *rol-6(su1006)*T animals. In contrast to what we observed with *rol-6(su1006)*T, exposure to *C. aquatica* dramatically increased rolling in *sqt-3(be3)*, *sqt-3(b238), dpy-10(cn64)* and *rol-1(sc2)* mutants (Figure 3A and B). Similar to what we observed with the *rol-6(e187)* mutants, rolling was not influenced by diet in the highly penetrant *rol-9(sc148)* mutants. In contrast to what we observed with *rol-6(su1006)*T animals, L1-arrest increased the penetrance of rolling in adult *sqt-3(be3), sqt-3(b238)*, and *dpy-10(cn64)* mutants (Figure 3A). All together, we noted a dichotomy in *rol-6(su1006)*T animals and other collagen mutants, conditions that reduced rolling in *rol-6(su1006)*T animals increased rolling in the other mutants. Although diet and arrest had opposing effects in *sqt-3* and *rol-6* mutants, *sqt-3(bc8)* mutant animals, like *rol-6(su1006)*T, displayed increased Hoechst staining when exposed to *C. aquatica* (Figure 3C). These data suggest that exposure to a *C. aquatica* diet, or to L1 arrest, does not produce a more robust cuticle resistant to aberrant collagen proteins, but rather causes a change in the composition or structure of the cuticle that differentially affects *sqt-3* and *rol-6* mutants.

**Figure 3.**
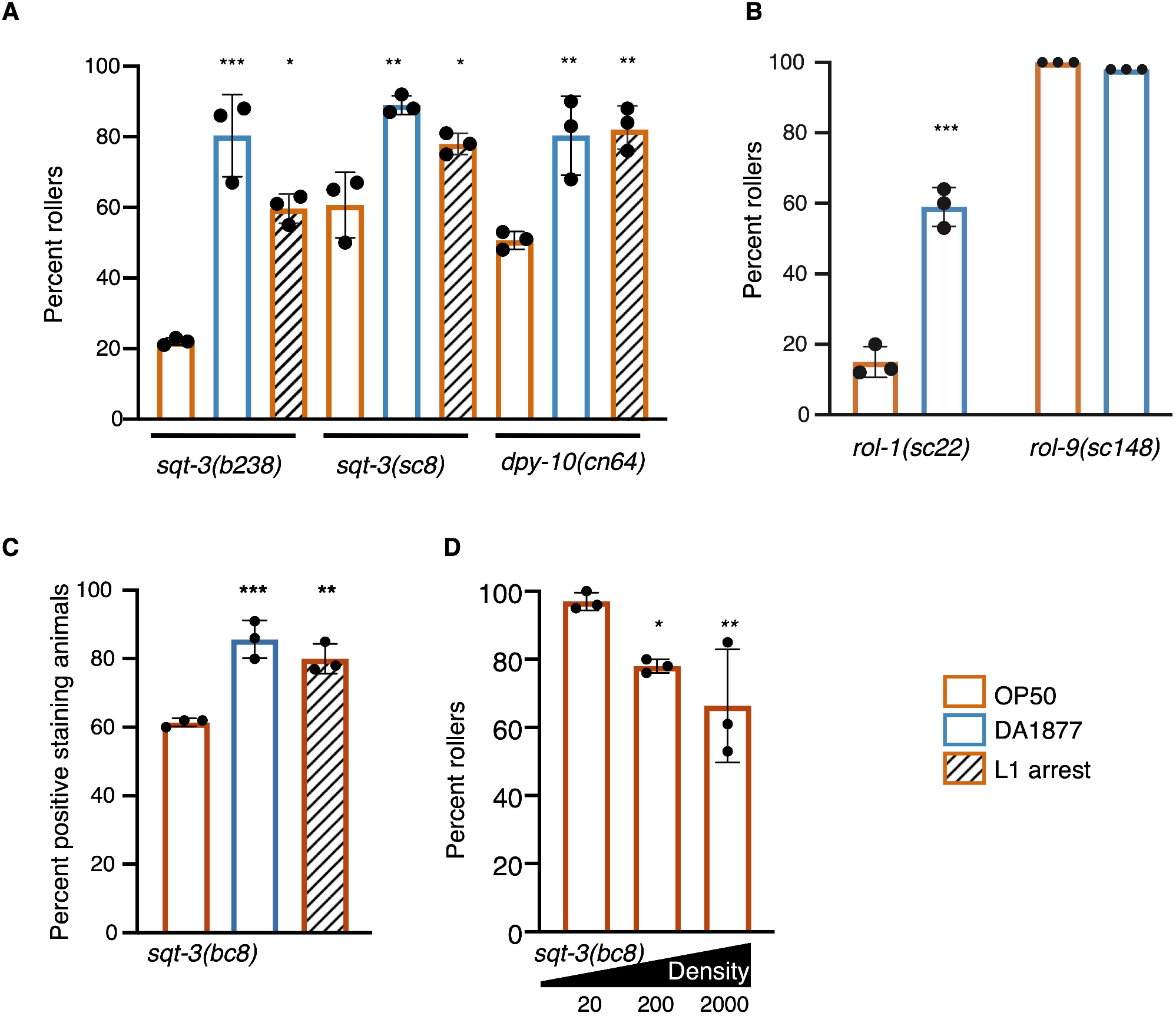
Environmental conditions affect the penetrance of rolling in many different mutants. A. Rolling in increased in *sqt-3* and *dpy-10* mutant adults that previously underwent L1 arrest or are fed *C. aquatica* (DA1877). B. Rolling in increased in response to *C. aquatica* in *rol-1*, but not *rol-9(sc148)* mutants that exhibit complete penetrance independent of diet. C. Percent *sqt-3(sc8)* animals with Hoechst stained nuclei. D. Rolling is decreased as population density increases in *sqt-3(bc8)* mutants. Numbers indicate size of population in a 60mm dish. Bars represent mean, plus and minus the standard deviation. Statistical analysis by one-way Anova with Tukey’s multiple comparison test for A,C and D, t-test for B. Asterisks indicate P values (*<0.05, **<0.005, ***<0.0005).

Given the differential effects on L1 arrest and *C. aquatica* diet on rolling in *rol-6(su1006)*T and *sqt-3(bc8)* animals, we tested the effects of population density on rolling in *sqt-3(bc8)* mutants. Population density increased the penetrance of rolling in *rol-6(su1006)*T animals. We asked whether density would affect *sqt-3(bc8)* in the same way it affected *rol-6(su1006)*T animals or whether the effect on penetrance would again be reversed. Similar to the other environmental exposures, high population density produced the opposite effect in *sqt-3(bc8)* mutants as in the *rol-6(su1006)*T animals (Figure 3D). These data are consistent with the idea that in response to environment, *C. elegans* alters its cuticle, relying more heavily on different collagens under different environmental conditions. For example, the post L1-arrest adult cuticle may have a decreased dependence on ROL-6 and an increased dependence on SQT-3 and DPY-10, explaining why arrest decreases rolling in *rol-6* mutants but increases rolling in *sqt-3* and *dpy-10* mutants.

### Collagen gene expression is regulated in response to environment

The finding that three very different environmental conditions produce opposing effects on rolling in *sqt-3* and *rol-6* mutants may indicate that a central regulatory mechanism dictates these effects in all three environments. One possible mechanism of this regulation is transcriptional regulation of collagen gene expression. We used available gene expression data to examine collagen gene expression in response to different bacterial diets or exposure to bacterial and fungal pathogens (Coolon *et al.* 2009; Engelmann *et al.* 2011; MacNeil *et al.* 2013). Collagen gene expression was altered in response to bacterial exposure and in fact, of 173 collagens, only 23 were not affected by any of these bacterial or fungal exposures. From this set, we noted that both *rol-6* and *sqt-3* are regulated in response to specific bacterial exposures (Supplemental Table S1). With this idea in mind, we selected a subset of collagens to measure gene expression in response to a diet of *C. aquatica* and to L1 arrest. We included a subset that were responsive to diet, some that have previously reported genetic interactions with *sqt-3* or *rol-6* and others, including *col-141* and *col-142*, for which regulatory mechanisms had been described.

Using nCounter assays, we measured mRNA expression of these collagens, and known regulators of collagen expression in wild-type adults exposed to *C. aquatica* or L1 arrest and compared these to control animals fed *E. coli* OP50 (Figure 4A). Because exposure to *C. aquatica* and L1 arrest had similar effects in all strains tested, we focused on collagens whose expression changed, relative to animals fed *E. coli* OP50, in response to both conditions. Strikingly, expression of *dpy-13*, *lon-3*, *rol-6*, and *sqt-1* was decreased in both conditions relative to control animals grown on *E. coli* OP50 (Figure 4A). Suppression of rolling in *rol-6(su1006)*T animals could be explained by downregulation of *rol-6* which would also decrease expression of the transgene. Further, rolling in *rol-6* neomorphic alleles is suppressed by a *lon-3* null allele and by knockdown of *sqt-1* or *dpy-13* (Nyström *et al.* 2002; Cai *et al.* 2011). We validated this finding in the *rol-6(su1006)*T animals, and found that, indeed, knockdown of *lon-3*, *sqt-1*, or *dpy-13* suppressed rolling. Decreased expression of these genes in response to *C. aquatica* or L1 arrest could explain the decreased penetrance of rolling observed in *rol-6(su1006)*T animals. However, knockdown of *lon-3* and *dpy-13* also reduces rolling in *sqt-3(bc8)* mutants (Figure 4C), and therefore do not explain enhancement of rolling in *sqt-3* mutants. Our analysis of collagen gene expression was not exhaustive and increased or decreased expression of another collagen that exacerbates rolling in *sqt-3(bc8)* animals, but not *rol-6 (su1006)*T animals may explain the differential effects observed in these mutants.

**Figure 4.**
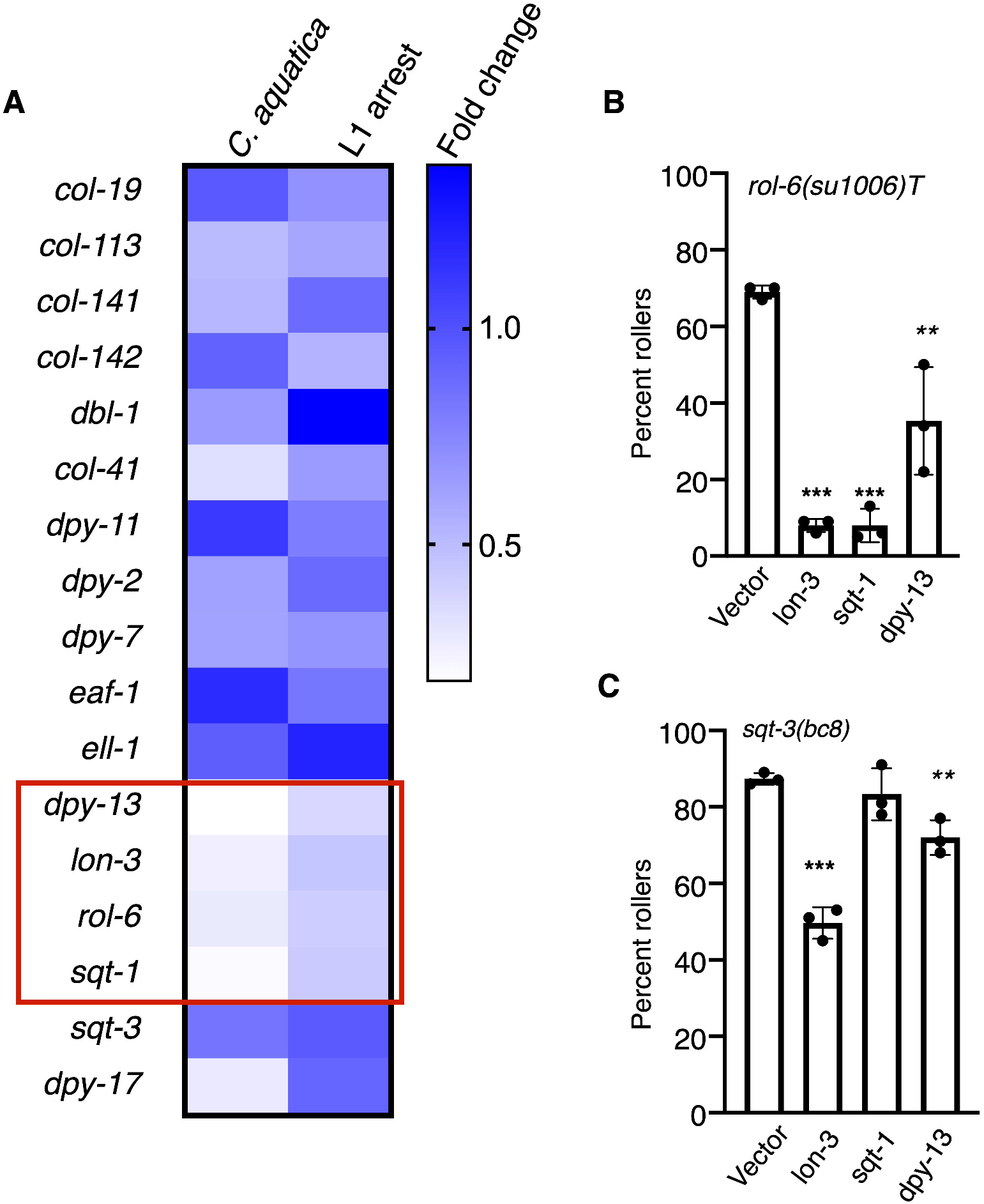
Environment influences expression of cuticular collagens. Expression of collagens, and regulators, was measured in wild type animals (N2) using nCounter technology. Of collagen genes tested only *dpy-13*, *lon-3*, *rol-6* and *sqt-1* (red box) produced statistically significant different changes in gene expression of 1.5-fold or more (p <0.05) in animals fed *C. aquatica* and in animals having experienced L1 arrest, relative to non-arrested animals fed *E. coli* OP50. Relative expression is shown, counts were normalized to expression in animals fed *E. coli* OP50. B. Knockdown of several collagen genes by feeding decreases rolling in *rol-6(su1006)*T animals (ERT60) and *sqt-3(bc8)* mutants. RNAi knockdown by feeding was used in a single generation. Each point represents a population of approximately 50 animals. Statistical analysis by one-way Anova with Tukey’s multiple comparison test. Asterisks indicate P values (*<0.05, **<0.005, ***<0.0005).

### ELT-3 regulates environmentally-induced changes in collagen gene expression

A decrease in the expression of *dpy-13*, *lon-3*, *sqt-1* and *rol-6* in response to two different conditions, L1 arrest or exposure to *C. aquatica*, suggests that these collagens may be targets of the same transcription factor (TF). Because conditions that increase rolling in *sqt-3(sc8)* mutants decrease rolling in *rol-6(su1006)*T, and *vice versa*, we searched for a TF whose loss would have opposing effects on rolling in these two genetic backgrounds. Using a cell-type specific dataset to identify TFs enriched in the hypodermis (Kaletsky *et al.* 2018) (Supplemental table 2), we knocked down each TF for which an RNAi clone was available and examined the penetrance of rolling in *rol-6(su1006)*T animals (Figure 5A). Two knockdowns produced significant changes in rolling, *elt-3*, which reduced rolling and *tbx-2*, which increased rolling (Figure 5A). We tested both knockdowns with a second *rol-6(su1006)*T strain (ERT60) and observed similar results (Figure 5B). To further extend our findings, we tested the effects of knocking down these two TFs on rolling in *sqt-3(sc8)* mutants. Knocking down these two genes produced opposite effects in *sqt-3(sc8)* and *rol-6(su1006)*T backgrounds, similar to what we observed with diet, density, and L1 arrest; *tbx-2* knockdown decreased rolling, whereas *elt-3* knockdown increased rolling (Figure 5C).

**Figure 5.**
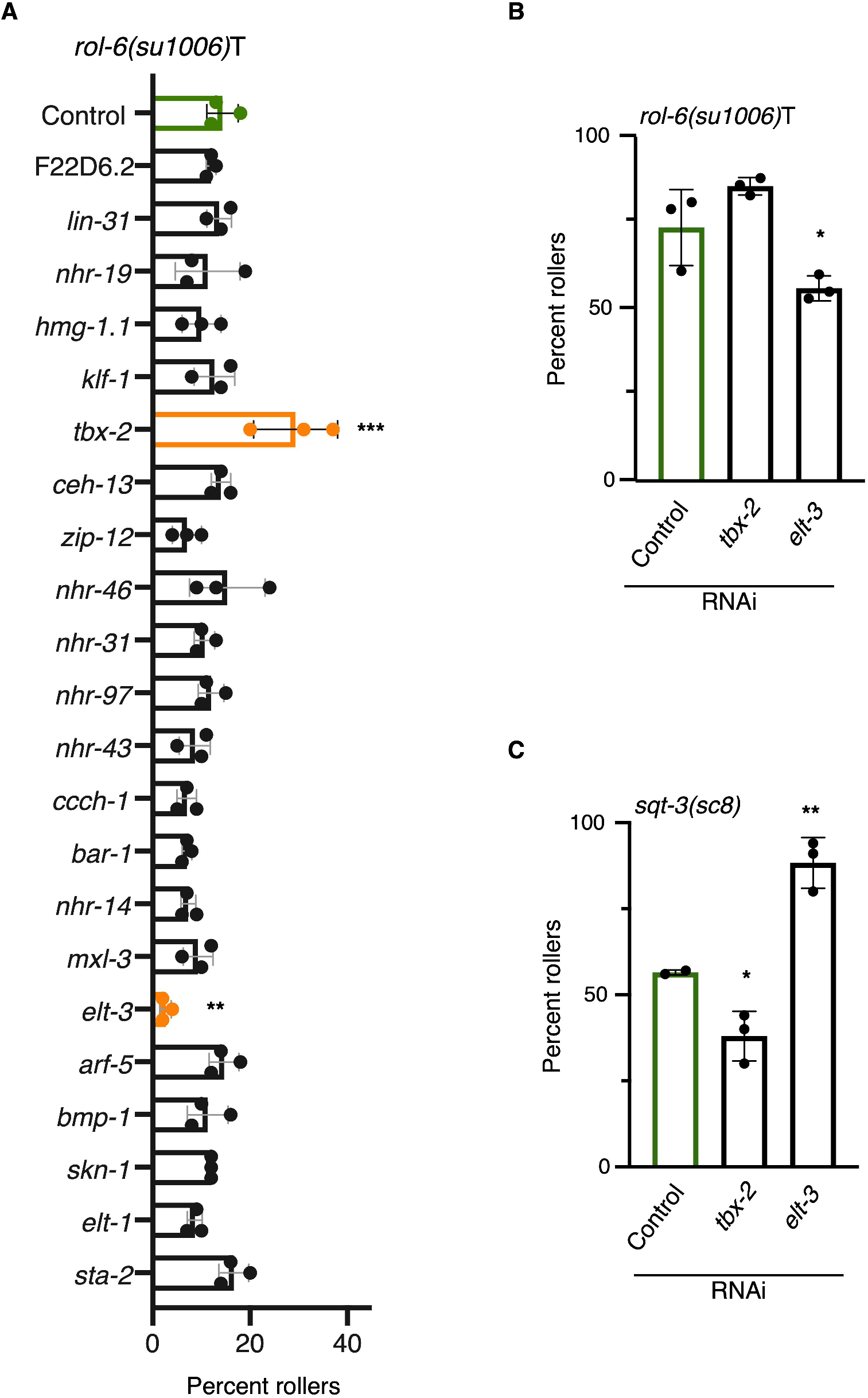
Identification of transcription factors mediating effects on rolling behaviour. A. RNAi mediated knockdown of individual transcription factors in *rol-6(su1006)*T animals. Knockdown of *tbx-2* and *elt-3* significantly increased, and decreased, respectively rolling in LMN40 *rol-6(su1006)*T animals. B. Knockdown of *tbx-2* and *elt-3* also significantly increased, and decreased (respectively) rolling in ERT60 *rol-6(su1006)*T animals. C. Knockdown of *tbx-2* and *elt-3* produce the opposite effect on rolling in *sqt-3(sc8)* mutants than in *rol-6(su1006)T* animals. Statistical analysis with one way Anova and Dunnett’s multiple comparison test. P-values * <0.05, **<0.005, ***<0.0005 ****<0.0001

To address whether ELT-3 directly regulates collagen gene expression, we too advantage of available ChIP-seq data (Gerstein *et al.* 2010) and examined the intergenic regions of *rol-6, lon-3, dpy-13* and *sqt-1* for the presence of ELT-3 binding sites. All four collagen genes have ELT-3 ChIP peaks in their intergenic regions (Supplemental Figure S1A), suggesting that ELT-3 may act as a direct transcriptional regulator of these collagens. In contrast, of the other collagens we measured, five out of 10, (including *sqt-3*) had significant ELT-3 ChIP peaks in their intergenic regions (Supplemental Figure S1B). To determine if ELT-3 was responsible for the downregulation of *rol-6*, *sqt-1*, *lon-3* and *dpy-13*, we measured the expression of these collagens following *elt-3* knockdown. Indeed, expression of all four genes was decreased when *elt-3* was knocked down (Figure 6A). These changes in gene expression are similar to what we observed in response to the *C. aquatica* diet or to L1 arrest (Figure 6B) suggesting that ELT-3 mediates changes in collagen gene expression in response to environmental stimuli.

**Figure 6.**
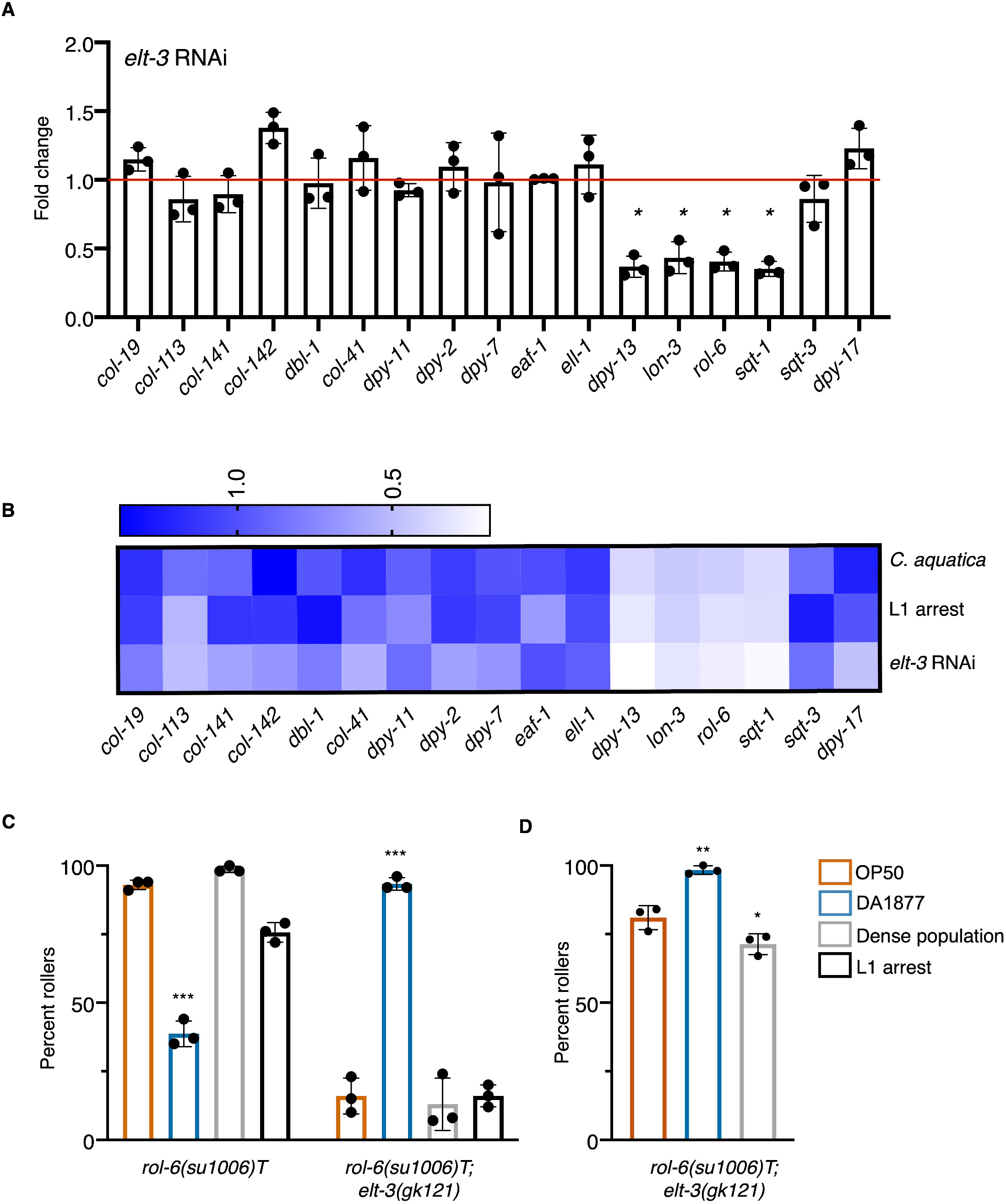
ELT-3 is a regulator of collagen gene expression. A. *elt-3* was knocked down in wild-type (N2) animals and mRNA expression of indicated collagens and collagen regulators was measured using nCounter assay. Expression is shown as fold change relative to treatment with empty vector control. Each point indicates a single replicate, bar represents the mean and error bars are standard deviation. P-values * <0.05 (t-test). B. Comparison of mean expression values for indicated treatments. L1 arrest and *C. aquatica* exposure are relative to animals grown on *E. coli* OP50. *elt-3* RNAi is shown relative to empty vector control in *E. coli* HT115. C. Analysis of rolling in ERT60 *rol-6(su1006)*T and *elt-3(gk121); rol-6(su1006)T* animals under well fed conditions. D. Rolling in response to indicated exposures in *elt-3(gk121); rol-6(su1006)T* in conditions that generated higher penetrance of rolling (populations that were routinely prepared from egg preps and allowed to starve before collecting eggs produced increased numbers of rollers).

To determine if *elt-3* was required for the environmental responses that influence rolling, we introduced the *elt-3(gk121)* mutation into *rol-6(su1006)*T animals (using ERT60) and examined the effect of diet, L1 arrest, and population density on rolling in these animals. Surprisingly, loss of *elt-3* did not inhibit the response to environment and its subsequent effect on rolling, but rather it reversed the effect of these factors on rolling. The *C. aquatica* diet increased rolling, while density decreased rolling (Figure 6C and 6D). The population remained sensitive to environmental factors. These effects would be consistent with a model where ELT-3 transcriptional targets are activated in response to *E. coli* OP50 or when populations are dense, but repressed when animals are fed *C. aquatica*. Our data suggest that ELT-3 plays a role in both activities.

## Discussion

The cuticle serves as a physical barrier that protects *C. elegans* from the outside environment. In different environments and in response to different stresses, there is likely an advantage to altering the composition of this collagen-rich structure. Our data support a model by which the GATA transcription factor ELT-3 mediates environmentally-induced changes in collagen gene expression that ultimately modify the cuticle. The effect of different environmental factors on rolling is reversed in *elt-3* mutants, suggesting that ELT-3 may function in both activating and repressing these genes. ELT-3 could accomplish this by functioning as an activator when animals are fed *E. coli* OP50 but functioning with a repressor protein when animals are fed *C. aquatica* (Figure 7). In this scenario, when ELT-3 is lost, expression of *elt-3* responsive genes would be decreased in normally activating conditions and increased in normally repressing conditions, generating the observed reversal of effects on rolling. Alternatively, ELT-3 may compete for binding to the same site as a repressor protein and the balance of the two proteins could dictate collagen levels.

**Figure 7.**
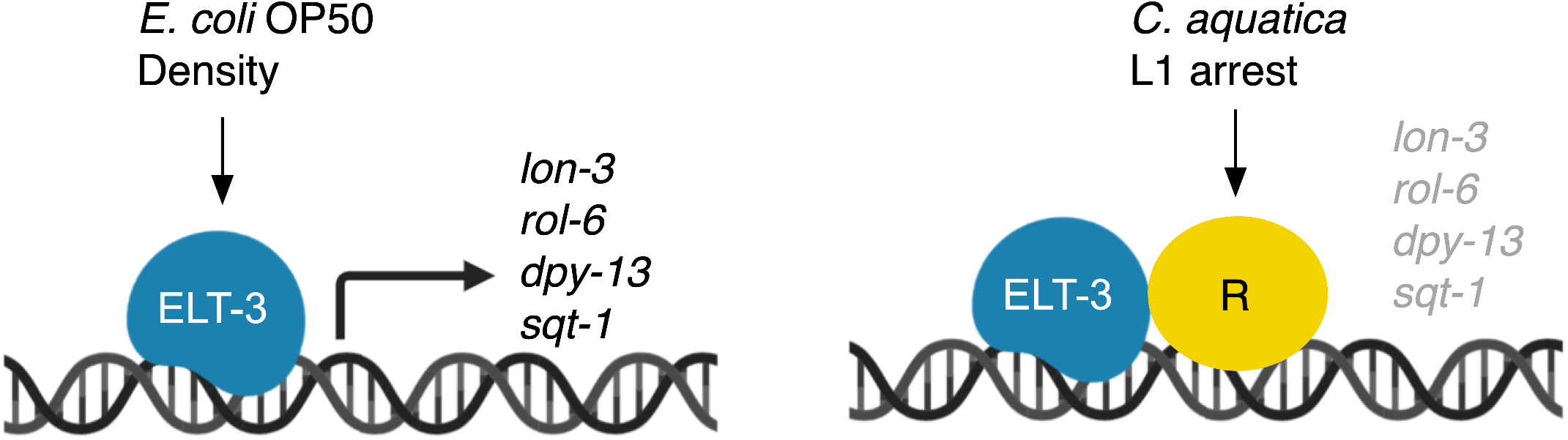
Expression of cuticular collagen genes is controlled by ELT-3. ELT-3 promotes expression of *dpy-13*, *lon-3*, *rol-6*, and *sqt-1* when animals are fed *E. coli* OP50 but contributes to the repression of these genes when animals are fed *C. aquatica* or following L1 arrest. This may occur through the recruitment of a repressor protein whose activity or recruitment to the promoter requires ELT-3.

The cuticle is replaced at each larval stage during development. Our findings that early life exposure, or exposure in the previous generation, can influence the assembly of the adult cuticle suggests that *C. elegans* use prior experience to tailor the cuticle later in life. This is consistent with the observation that the penetrance of rolling in *sqt-2(sc64)* mutants is influenced by developmental events, *sqt-2(sc64)* mutants derived from L3 animals are Sqt (homozygotes are Dpy and heterozygotes are Rol), whereas homozygotes derived from dauers are rollers (Cox *et al.* 1980). Changes in the presentation and penetrance of these defects may similarly occur through the modification of the composition of the cuticle, resulting in the aberrant SQT-2 protein having a greater impact in animals that have undergone dauer arrest. The idea that *C. elegans* modulates collagen expression in response to environment is also consistent with the finding that in the L2 stage preceding dauer (L2d), *rol-6* expression is not observed, whereas it is observed in L2 animals not undergoing dauer (Park and Kramer 1994).

How SQT-3 and ROL-6 proteins interact in the cuticle is unclear. While a balance between ROL-6 dependence and SQT-3 dependence is an attractive hypothesis based on the environmental effects observed in this study, it is unlikely that these proteins substitute for one another. SQT-3 is predicted to encode a transmembrane collagen, while ROL-6 is predicted to encode a globular collagen (Teuscher *et al.* 2019). Secondly, redundancy in the function of these two proteins would not be consistent with the fact that knockdown of *sqt-3* can suppress rolling in *rol-6* mutants (Cai *et al.* 2011). One proposed explanation for this suppression is that SQT-3 and ROL-6 are part of the same structure, along with other suppressing collagens, and that in the absence of one of these proteins, collagen fibrils containing aberrant proteins are decreased. However, the effects that we observe in these two mutants in response to environment cannot be explained by this simple model.

We observed decreased expression of *dpy-13, rol-6, lon-3,* and *sqt-1* in conditions that suppress rolling in *rol-6(su1006)*T animals but enhance rolling in *sqt-3(bc8)* animals. While decreased expression of these genes is consistent with suppression of *rol-6(su1006)T*, it does not explain the increased penetrance of rolling in *sqt-3* mutants. Our analysis of collagen gene expression was not comprehensive and it seems likely that the downregulation of these four collagen genes is matched with the upregulation of other compensatory collagen genes that fill these roles. Additional changes in the expression of collagens, collagen interacting proteins, collagen modifying enzymes, or factors involved in assembly or secretion of collagens likely explain increased rolling in *sqt-3* mutants. We did not observe increased expression of *sqt-3*, which may have provided the most obvious explanation for the effects observed. However, the timing or duration of expression, or changes to the SQT-3 protein or its export from the ER could increase the penetrance of rolling in *sqt-3* mutants and may explain the observed effects.

ELT-3 has been implicated in both stress response and damage response in the cuticle (Pujol *et al.* 2008; Hu *et al.* 2017; Dodd *et al.* 2018). Disruption of annular furrows in the cuticle increases accumulation of osmolytes and activates specific stress response pathways that require ELT-3 (Dodd *et al.* 2018). ELT-3 regulation of collagens may also play a role in recovery from cuticle damage and pathogen resistance and as such, inability to regulate collagen gene expression in response to stress may contribute to the observed decreased resistance of *elt-3* mutants to the fungus, *Dreshmeria conispora*, a pathogen that adheres to the *C. elegans* cuticle (Pujol *et al.* 2008). ELT-3 may promote the production of a cuticle that is more resistant to infection by this pathogen in much the same way *col-92* overexpression increases resistance to *Bacillus thuringiensis* infection (Iatsenko *et al.* 2013). In addition to the regulation of *rol-1*, *sqt-1*, *lon-3* and *dpy-13* described here, ELT-3 is also reported to regulate the expression of *col-144* (Budovskaya *et al.* 2008)*, col-41* (Yin *et al.* 2015)*, col-34, and dpy-7* (Yin *et al.* 2015), suggesting that these additional collagens could also be modulated in response to environmental factors.

Modifying the cuticle may be most easily accomplished by transcriptional co-regulation of collagens that interact to ensure proper assembly of cuticle structures. In response to the *E. coli* diet, ELT-3 would therefore function to generate a cuticle richer in ROL-1, SQT-1, LON-3 and DPY-13 collagens. Transcriptional co-regulation of additional groups of collagens likely functions to generate cuticles optimized to specific developmental stages or environmental conditions. For example, NHR-23 regulates expression of *dpy-2*, *dpy-3*, *dpy-7*, *dpy-8*, *dpy-10* and *dpy-5* (Kouns *et al.* 2011) and LIN-29 is required for the expression of L4-specific collagens *col-38*, *col-49*, *col-63* and *col-138* (Abete-Luzi and Eisenmann 2018). In these cases, TFs function to ensure the production of stage-specific cuticles. Additional transcriptional regulators of collagen groups have been identified, including EAF2 and ELL-1 that regulate expression of *dpy-3*, *dpy-13*, and *sqt-3* (Cai *et al.* 2011), and BAR-1, that regulates *bli-1*, *col-38*, *col-49* and *col-71* (Jackson *et al.* 2014). TGF-β signaling regulates expression of *col-141* and *col-142* (Madaan *et al.* 2018). Intriguingly, TGF-β signaling has been implicated in environmental response, raising the possibility that modification of the cuticle may also be integrated into these responses.

How vitamin B12 promotes changes to the cuticle is not clear. However, the observation that when Vitamin B12 is depleted, infrequent Dpy animals appear in the population, suggests that B12 normally contributes to cuticle development (Bito *et al.* 2013). One possible explanation for the connection between vitamin B12 and ELT-3 is oxidative stress. Depletion of vitamin B12 increases oxidative stress in *C. elegans* (Bito *et al.* 2017) and although the *E. coli* OP50 diet is not devoid of vitamin B12, the low levels of vitamin B12 in the *E. coli* diet, relative to the *C. aquatica* diet, may promote higher levels of detoxification responses. ELT-3 has been identified as a mediator of these responses (Hu *et al.* 2017), which may suggest the link between vitamin B12 availability and ELT-3 activation is stress response. If true, ELT-3 would couple the induction of detoxification response with changes in the cuticle. Based on our observations with Hoechst staining, an *E. coli* OP50 diet, when ELT-3 targets are more highly expressed, results in a less permeable cuticle. Decreasing the permeability of the cuticle may provide additional protection from conditions that activate detoxification responses.

Our model suggests an unknown repressor protein participates in the environmentally induced regulation of cuticular collagens. One candidate for this repressor is TBX-2, identified in our RNAi screen as having an activity opposite to that of ELT-3. TBX-2 is the sole *C. elegans* homolog of Tbx2 subfamily of T-box factors. In humans, TBX2 can act as a repressor or an activator, however in *C. elegans*, it has only been described as a repressor. Intriguingly, in mammalian cells, TBX2 is proposed to act as a regulator of the collagen COL1A2. Stable expression of TBX2 reduces expression of COL1A2 in fibroblast cell lines (Teng *et al.* 2007). However, in osteoblast cells, TBX2 expression has the opposite effect on COL1A2 expression. Intriguingly, COL1A2 is also regulated by a GATA factor, GATA4, suggesting the potential for the roles of TBX-2 and ELT-3 to be conserved in mammalian cells.

Altogether, we find that diet, population density, early life starvation and parental environment impact cuticle formation. Transcriptional regulation of cuticular collagens provides one mechanism by which *C. elegans* adapts to the surrounding environment. We identified ELT-3 as one factor involved in this regulation, however, many other regulators of collagen gene expression have been identified. These, and other regulators, likely function to tailor the cuticle to specific environmental conditions.

## Methods

### *C. elegans* propagation

*C. elegans* were propagated by standard methods (Stiernagle 2006). Soy peptone was used in Nematode growth media (NGM). The following strains used were obtained from the CGC: NR222 *rde-1(ne219)*; *kzIs9 [pKK1260(lin-26p∷nls∷GFP) + pKK1253(lin-26p∷rde-1) + pRF6(rol-6(su1006)]*, VP303 *r<u>de-1</u>(<u>ne219)</u>*; *kbIs7[nhx-2p∷rde-1 + rol-6(su1006)](Espelt et al. 2005)*, FK181 *ksIs2[pdaf-7∷GFP + rol-6(su1006)]* (Murakami *et al.* 2001), MS438 *irIs25*[*elt-2*∷NLS∷GFP∷lacZ + *rol-6(su1006)*], CB187 *rol-6(e187) (Kramer and Johnson 1993)*, BE22 *rol-1(sc22)*, BE148 *rol-9(sc148) (Bergmann et al. 1998)*, BE8 *sqt-3(bc8)*, *elt-3(gk121)*. LMN40 was generated by outcrossing VP303 to N2 to remove the *rde-1(ne219)* mutation. *elt-3(gk121)* was crossed to ERT60 to generate *elt-3* mutants with the *rol-6(su1006)* transgene.

### Scoring rolling animals

Rolling was scored in adult animals for all strains. Plates were tapped to promote movement and animals rolling in a circle or rolling on their longitudinal axis were counted as rollers. Three replicate plates were scored for each condition. For most assays, 50-100 eggs were added to each prepared plate and animals were scored as adults. Density experiments were plated as indicated in Figures 1 and 2. NR222 was scored at 25°C because they did not produce appreciable numbers of rollers at 20°C, all other animals were scored, and maintained, at 20°C. Significance of rolling effects was measured with one way Anova and Dunnett’s multiple comparison test in Graphpad Prism.

### Hoechst staining

Hoechst staining was performed as described previously (Moribe *et al.* 2004). Briefly, adult animals were washed off plates with M9, and washed to remove bacteria before addition of 1μg/mL Hoechst 33258 (Sigma). Animals were placed on a rocker and incubated at room temperature for 20 minutes. Following incubation, animals were washed twice with M9 buffer, treated with 1mM levamisole and examined on a Nikon Eclipse II microscope.

### L1 arrest and Starvation

Eggs were collected by hypochlorite bleaching, pelleted by centrifugation and washed 3 times in M9 buffer. Collected eggs were incubated in M9 buffer for 24 hours with rocking to allow for hatching, synchronization, and starvation. 50-100 L1s were added to prepared plates. Animals were scored in the adult stage.

### Density

Eggs were plated at indicated density from a single pool of eggs. Additional *E. coli* OP50 was added daily to ensure worms did not starve. An overnight culture of *E. coli* OP50 was pelleted by centrifugation and 1 ml of the concentrated bacterial suspension was added to each plate. For plates with more than 200 worms, one quarter of each plate was counted.

### nCounter measurements

N2 Eggs were prepared by hypochlorite treatment, plates were seeded and animals developed at 20C. Animals were collected when the population was a mix of late L4 and early adult animals, washed 3 times in M9 and frozen at −80°C in Trizol (Sigma) before RNA preparation. RNA was extracted according to manufacturer’s protocol and further purified using NEB RNA columns. nCounter probes (nanoString Technologies) were designed by nanostring, synthesized by Integrated DNA Technologies (IDT) and assays were performed by Mobix, McMaster University, Hamilton, Ontario. nSolver software was used for analysis. The geometric mean of positive control probes was used for in-lanes normalization. Counts were normalized across samples using the geometric mean of *pmp-3*, Y45F10D.4 and *cdc-42* counts (Hoogewijs *et al.* 2008). Statistically significant changes were identified by t-test using nSolver software. Probes used are listed in supplemental table 3. Additional probes that were used but not reported were excluded because counts were close to, or below, negative threshold levels in multiple samples (as determined by negative probe counts).

### Vitamin B12 treatment

Methylcobalamin (Vitamin B12) (Sigma) was diluted in water to 5 mg/mL, filter sterilized, and stored at −20°C until use. Stocks were diluted in sterile water and 40 μL of diluted B12 solution was added to the surface of a 60 mm plate after growth of the *E. coli* OP50 lawn at doses indicated in Figure 1 and allowed to dry. Worms were added to plates immediately.

### RNAi

The *mmcm-1* clone was obtained from the ORFeome RNAi library (Rual *et al.* 2004). RNAi clones for *dpy-13*, *lon-3* and *sqt-1* were generated by PCR, inserted into L4440 and transformed into *E. coli* HT115. Small, unique regions were selected to reduce off-target effects (Table S3). All other RNAi clones were obtained from the Ahringer RNAi library (Kamath *et al.* 2003). RNAi clones were inoculated into LB supplemented with 60mg/mL ampicillin. 50 ◻L of overnight culture was used to reinoculate 3 mL of LB supplemented with ampicillin and grown for six hours at 37°C before being used to seed RNAi plates. RNAi plates were made by adding 1mM IPTG and 25 μg/ml carbenicillin to NGM made with soy peptone. RNAi cultures were seeded and grown overnight at 37°C. Positive acting clones were sequenced to verify their identity. Three replicates were counted for each knockdown.

## Data Availability

Strains and plasmids are available upon request. The authors confirm that all data necessary for confirming the conclusions presented are within the article, figures, and tables.

## Acknowledgments

We thank S. Taubert and T. Kubiseski for comments on the manuscript. Some strains were provided by the CGC, which is funded by NIH office of Research Infrastructure Programs (P40 OD0104440). We gratefully acknowledge the support of the Natural Sciences and Engineering Research Council of Canada (NSERC), [RGPIN-2016-06339].

---

Supplemental Figure 1. A. ChIP-seq data of ELT-3 binding to indicated collagen gene promoters. B. Summary of ELT-3 binding in the intergenic regions of indicated collagens (summarized from modEncode ChIP-seq data).

Table S1.

Collagen gene expression in response to bacterial exposures. Data summarized from (Coolon *et al.* 2009; Engelmann *et al.* 2011; MacNeil *et al.* 2013). Genes that were upregulated are shown in orange, downregulated in yellow. Classification of collagen genes as defined in (Teuscher *et al.* 2019). Collagens highlighted in red were selected for measuring in nCounter assays.

Table S2

Hypodermally enriched transcription factors. The overlap between TF2.0 transcription factors and genes enriched in the hypodermis (Kaletsky *et al.* 2018) was used to create a list of candidate regulators.

Table S3.

Nanostring probes and sequences used in nCounter analysis. RNAi target sequences.

